# Non-differentiable activity in the brain

**DOI:** 10.1101/2023.06.12.544700

**Authors:** Yasuhiro Tsubo, Shigeru Shinomoto

## Abstract

Spike rasters of multiple neurons show vertical stripes, indicating that neurons exhibit synchronous activity in the brain. We seek to determine whether these coherent dynamics are caused by smooth brainwave activity or by something else. By analyzing biological data, we find that their cross-correlograms exhibit not only slow undulation but also a cusp at the origin, in addition to possible signs of monosynaptic connectivity. Here we show that undulation emerges if neurons are subject to smooth brainwave oscillations while a cusp results from non-differentiable fluctuations. While modern analysis methods have achieved good connectivity estimation by adapting the models to slow undulation, they still make false inferences due to the cusp. We devise a new analysis method that may solve both problems. We also demonstrate that oscillations and non-differentiable fluctuations may emerge in simulations of large-scale neural networks.

## INTRODUCTION

Technological advancements have made it possible to record signals from multiple neurons for hours or days [1– 5]. The number of recorded neurons has doubled every seven years as with Moore’s law for semiconductor integrated circuits [6]. This number now exceeds several thousand even for detecting electrical signals with sub-millisecond resolution [7–9]. This unprecedented increase in the amount of data is expected to change our fundamental understanding of neuronal functions in the brain, just as semiconductor technologies have changed our daily life.

There have been various proposals to use multiple spike signals, particularly for analyzing inter-neuronal connections [10–16]. It all started more than a half-century ago when Perkel, Gerstein, and Moore proposed to estimate the monosynaptic connectivity by capturing the interdependence between neuronal firings. This was done by detecting the significant deviations from a uniform distribution in a cross-correlation histogram a.k.a. a crosscorrelogram (CC) of spike trains recorded from a pair of neurons [17]. Nevertheless, the classical analysis methods make many false inferences because *in vivo* neuronal firings are originally correlated due to background activity even if a pair of neurons are not monosynaptically connected. There have been many attempts to improve the estimation [18–29], most of which involved the adaptation of models to variations that appeared in the CCs *ex post facto* by regarding them as unwelcome disturbers, but without determining the underlying causes.

Neuroscientists have seen many CCs exhibiting a sharp peak at the origin and presumed that such a peak may have been caused by common inputs from locally connected neurons. Here we analyze a large number of biological CCs and explore their possible causes. We conclude that CCs may exhibit both smooth undulation and a sharp peak (or non-smooth cusp) even if there are no specific structures of local connections. Namely, a CC exhibits smooth undulation if a pair of neurons are subjected to smooth oscillations, while it exhibits a cusp if there are external fluctuations that are steep enough to be expressed as “non-differentiable.”

While modern analysis methods have achieved a reasonable estimation of monosynaptic connectivity by adapting the models to slow undulation, they still make false inferences when a CC exhibits a sharp cusp at the origin. Accordingly, we revise the analysis method GLMCC developed by one of the authors [28] into a new method “ShinGLMCC” by allowing the model to adapt to not only undulation but also a cusp.

To test the reliability of analysis methods, we perform simulations of a large-scale network of neurons interacting with fixed synaptic connectivity. We find that the network may exhibit non-differentiable activity similar to what we have observed in biological spike trains. Theoreticians have known that coherent synchronous activity may emerge not only in neural networks [30–37] but also in various self-excitation processes [38]. While theoreticians have discussed possible functions for coherent dynamics in neuronal information processing, they have not paid much attention to the sharp temporal profile of the emergent fluctuating activity. The sharp fluctuations occurring in neuronal networks may be a basis for complex information processing.

We also devise fast algorithms for both GLMCC and ShinGLMCC and provide a user-friendly software package so that researchers can obtain a neuronal connection diagram and individual analysis details for all neuron pairs from a large number of recorded spike trains.

## RESULTS

### Cross-correlograms (CCs) of biological data

The CC is a basic device to display the potential interdependence of two units. This can be constructed by simply superposing the spike times of one train measured relative to every spike of another train (Fig. 1a). Here we demonstrate a variety of CCs obtained from publicly sourced data of spike trains recorded in parallel from wide brain regions of a freely moving rat using a silicon probe called Neuropixels [39].

**FIG. 1.**
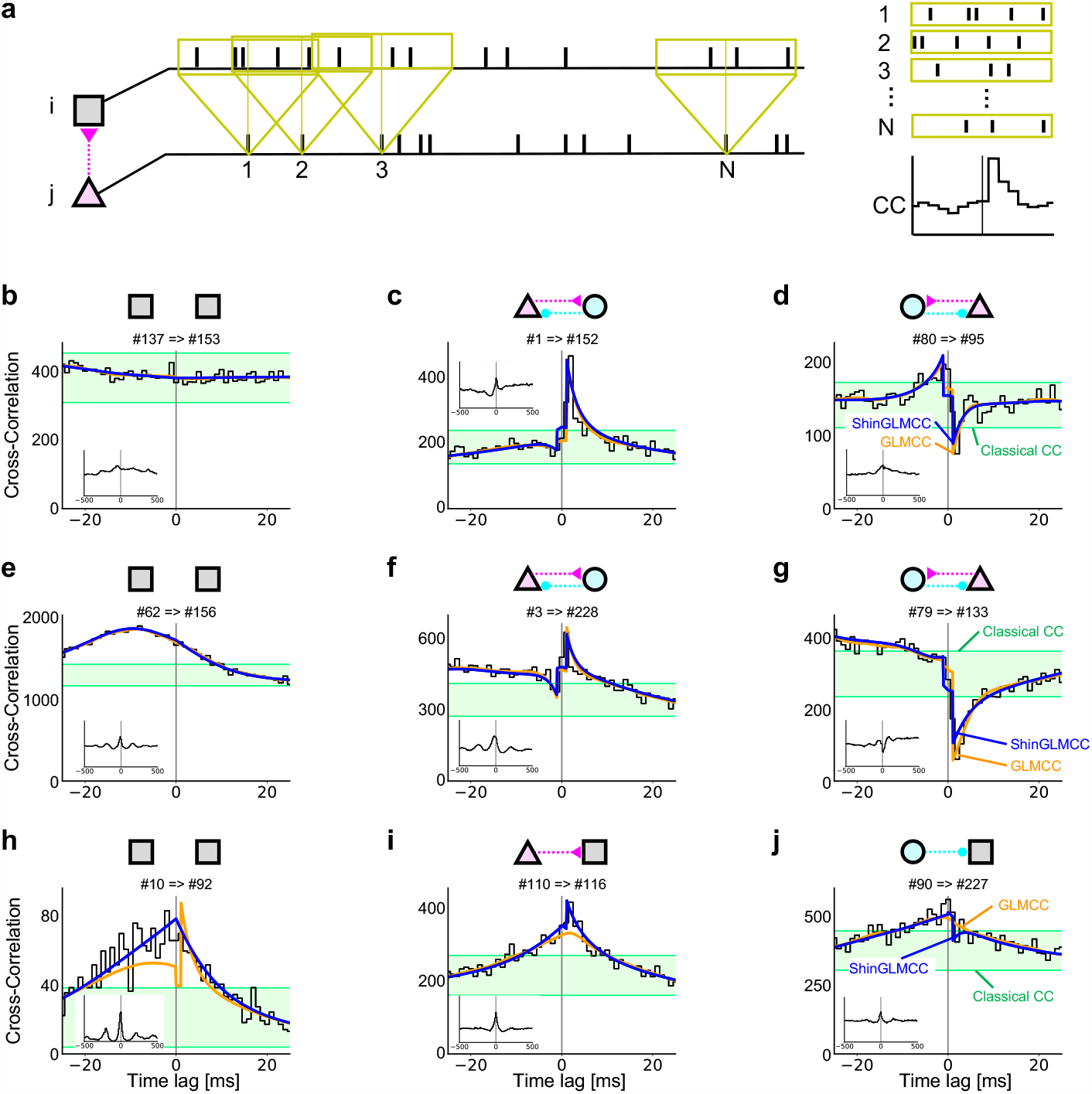
A variety of cross-correlograms (CCs) obtained from biological spike trains. **a** Process of constructing a CC from a pair of spike trains. **b** A flat CC. **c** and **d** A hump and a dip at several milliseconds right side from the origin. **e** A CC exhibits a large undulation. **f** and **g** A hump and a dip immersed in large undulating variations. **h** A CC exhibits a cusp at the origin. **i** and **j** A hump and a dip immersed in a cusp. Green belts represent a confidence interval suggested by the Classical CC analysis. Orange and blue lines represent inferences of GLMCC and ShinGLMCC, respectively. Schematic diagrams in the above represent putative connections determined by ShinGLMCC. Triangles and circles represent putative excitatory and inhibitory neurons, while squares are for those with undetermined characteristics.

#### Flat distribution in CCs

If a pair of neurons has generated spikes independently of each other, their CC is expected to be flat apart from statistical fluctuations that occur when counting spikes in each bin. A large fraction of neuron pairs in the cortex display flat CCs, implying that synaptic connections between them may be either absent or insignificant, as shown in Fig. 1b.

If synaptic connections exist between neurons, they will induce statistical interdependence between two spike trains, and their CC is expected to display either a hump for excitatory monosynaptic connectivity (Fig. 1c) or a dip for inhibitory connectivity (Fig. 1d) several milliseconds from the origin. A “Classical CC analysis” seeks to detect a significant deviation from the flat distribution based on the stochastic point process theory [17, 40]. The confidence intervals representing natural statistical fluctuations are depicted as green belts in Fig. 1.

#### Smooth undulation in CCs

While Classical CC analysis gives plausible inferences for typical cases (Figs. 1b–d), it does fail in many cases, particularly when there are large undulating variations that exceed the assigned confidence interval as in Figs. 1e–g. There have been many efforts to reduce the influence of such large variations [18–27]. Most of them attempted to smooth out such large variations by regarding them as unwelcome disturbers. For instance, one of the authors developed a method called GLMCC, in which a generalized linear model adapts a slowly fluctuating function to existing variations in the CC, and accordingly, reduces false inferences [28]. The analysis method was further revised by introducing the likelihood ratio test into the determination of connections [29]. The slowly fluctuating functions and monosynaptic impacts determined by GLMCC are depicted with orange lines in Fig. 1. Large undulations are well represented by GLMCC, particularly in Figs. 1e–g.

#### A cusp in CCs

By analyzing a large number of biological spike trains, however, we have newly found other difficult cases that even the GLMCC makes dubious inferences for. This occurs particularly when a CC exhibits a seemingly non-smooth cusp at the origin as represented in Figs. 1h–j. In these cases, GLMCC (plotted in orange lines) does not seem successful in representing large variations in individual CCs.

In the present study, we develop ShinGLMCC from GLMCC so that the adaptation function can bend at the origin as shown in blue lines in Figs. 1h–j. Before developing the analysis method, we consider the possible cause for the cusp that appears in CCs in the following section.

### Models for the variations in CCs

To tackle the problem that accompanies both the undulation and the cusp in CCs, we begin by exploring their possible causes using mathematical models for spike generation (METHODS).

#### Producing smooth undulations in a CC

Before addressing the puzzle of the cusp, let us first consider the appearance of smooth undulation in a CC. Smooth undulation may be produced by adding smooth oscillations in the firing rates of a pair of neurons, mimicking the situations that biological neurons are influenced by (smooth) brainwave oscillations (METHODS). The basic frequency of the undulation in a CC is identical to the frequency of the rate modulation. The undulation may have a peak at the CC’s origin if two neurons are influenced in phase. The undulation may exhibit damping if the frequency components of the rate modulation are distributed (Fig. 2a). The peak of the undulation in a CC may be shifted if a pair of neurons receive oscillatory modulation with a certain time lag (Fig. 2b). CCs constructed from a given data set exhibit fluctuations in counting spikes as long as the spike trains are of finite length. Nevertheless, such counting fluctuations can be minimized if spike trains are sufficiently long because the number of spike counts in each bin of the CC histogram increases with the length of spike trains.

**FIG. 2.**
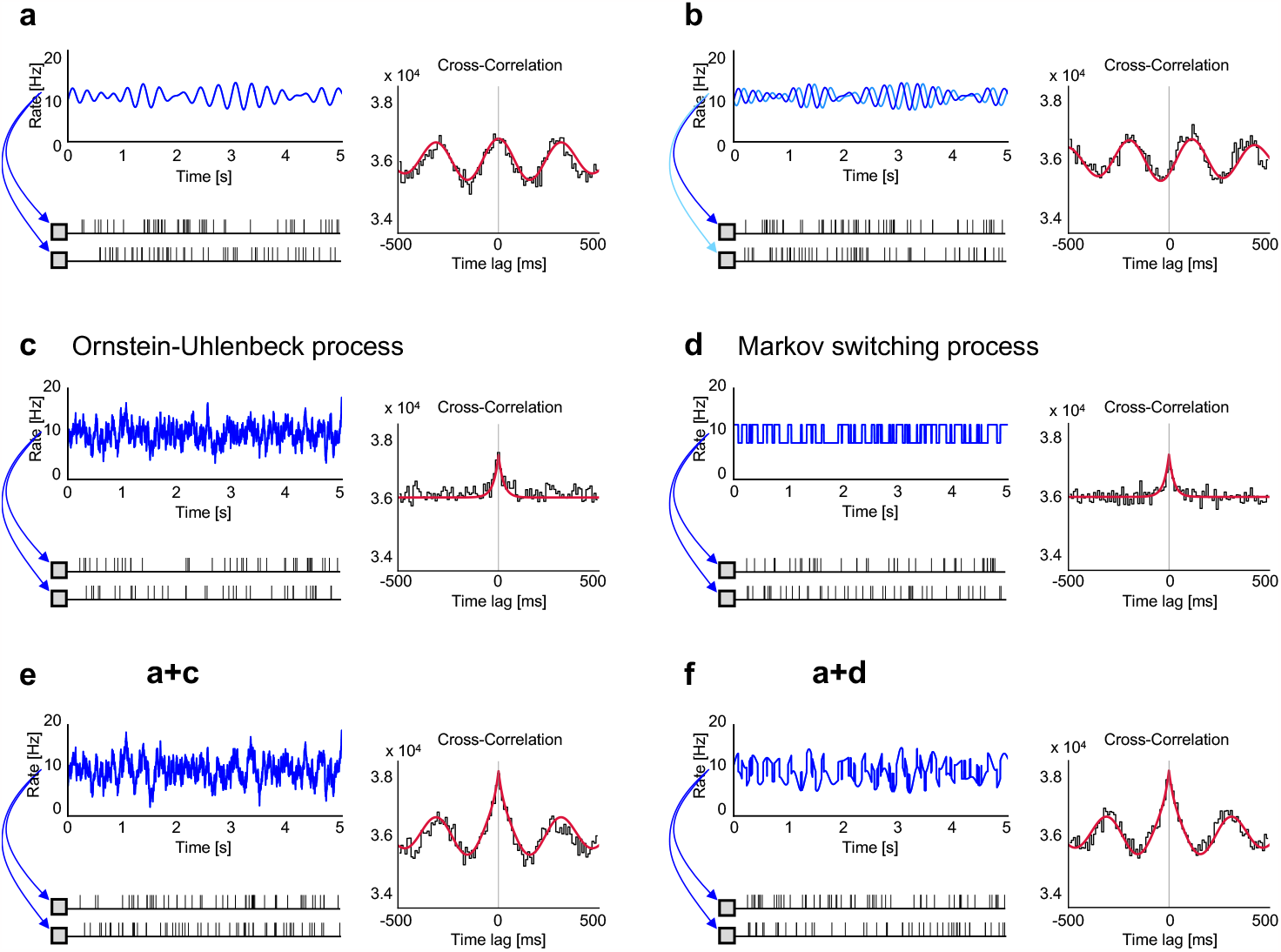
Synthetic spike trains producing the smooth undulation and a cusp in CCs. **a** Neurons are commonly modulated with smooth oscillations. **b** The case of smooth modulation with a time lag. **c** Neurons are commonly modulated with rates that are non-differentiable everywhere, according to the Ornstein-Uhlenbeck process. **d** Neurons whose firing rates switch between high and low values randomly in time, according to the Markov switching process. **e** Neurons are modulated by oscillations and non-differentiable fluctuations (**a** and **c**). **f** Neurons are switching between high and low oscillatory states (**a** and **d**). Red lines are analytically obtained CCs for individual models.

#### Producing a cusp in a CC

While a CC can show smooth undulation due to background oscillations as described above, it will never show a (non-smooth) cusp as long as external influences are smooth. A necessary and sufficient condition for having a smooth correlation function is that the original rate processes are “mean square differentiable” [41]. Accordingly, a pair of neurons that show a cusp in the CC must have been modulated by non-differentiable activity.

We have explored possible spike-generation processes such that the CC shows a cusp at the origin (METHODS). One possible situation is that neurons generate spikes with time-dependent firing rates that are non-differentiable everywhere, as exemplified by the “Ornstein-Uhlenbeck process” (Fig. 2c) [42–45].

Note that this model process assumes that two neurons are not monosynaptically connected, because spikes of one neuron have never influenced the spike generation of another neuron. Nevertheless, their firings are not independent, because they are influenced by identical background fluctuations. The model simulation mimics the situation that neurons are influenced by common fluctuations that are non-differentiable everywhere.

We also found another situation in which the CC exhibits a cusp. Namely, two neurons generate spikes according to firing rates that switch between high and low values randomly according to the “Markov switching process” (Fig. 2d) [45–49]. While the underlying rate is not non-differentiable everywhere, it exhibits (more drastic) discontinuous changes at a finite rate. Two neurons are not monosynaptically connected either, because spikes of one neuron do not influence the firing of another neuron.

Different rate processes such as the Ornstein-Uhlenbeck process and Markov switching process may end up with analytically identical CCs exhibiting a cusp at the origin. We cannot identify a time profile of the original rate process accurately from the recorded spike trains because spikes are a mere random realization of a stochastic process. Nevertheless, we can obtain a sharp profile of the CC from sufficiently long spike sequences because the number of spikes in histogram bins can be increased with the recording duration, such that the relative influence of the spike count fluctuations can be made sufficiently small. The cusp at the origin of a CC may remain even sharply if we can collect longer time series.

Figures 2e and 2f depict examples of a pair of spike trains derived from the rate process in which smooth oscillation of Fig. 2a and non-differentiable fluctuations of Figs. 2c and 2d are mixed. In this case, the CC exhibits not only smooth undulation but also a cusp at the origin, as seen in the biological data in Figs. 1h–j.

### Population activity of biological data

To observe the non-differentiable feature of the background activity, which may have caused a cusp at the origin of pairwise CCs, we summed a large number of spike trains into a single spike train, expecting the background activity to appear in this single spike train’s firing rate. A time histogram of an entire spike train of 242 units is depicted in Fig. 3a. Slow oscillations and seemingly non-differentiable activity coexist in an entire firing rate. Raster diagrams of individual spike trains are shown in Fig. 3b, in which we observe a striped pattern, indicating that a subset of neurons exhibits seemingly sharp fluctuations in their firing rates on the order of 10 −100 ms. To see the degree of temporal variability of the original data, we show a time histogram and rasters of shuffled spike trains on the right side of Figure 3a as a reference.

**FIG. 3.**
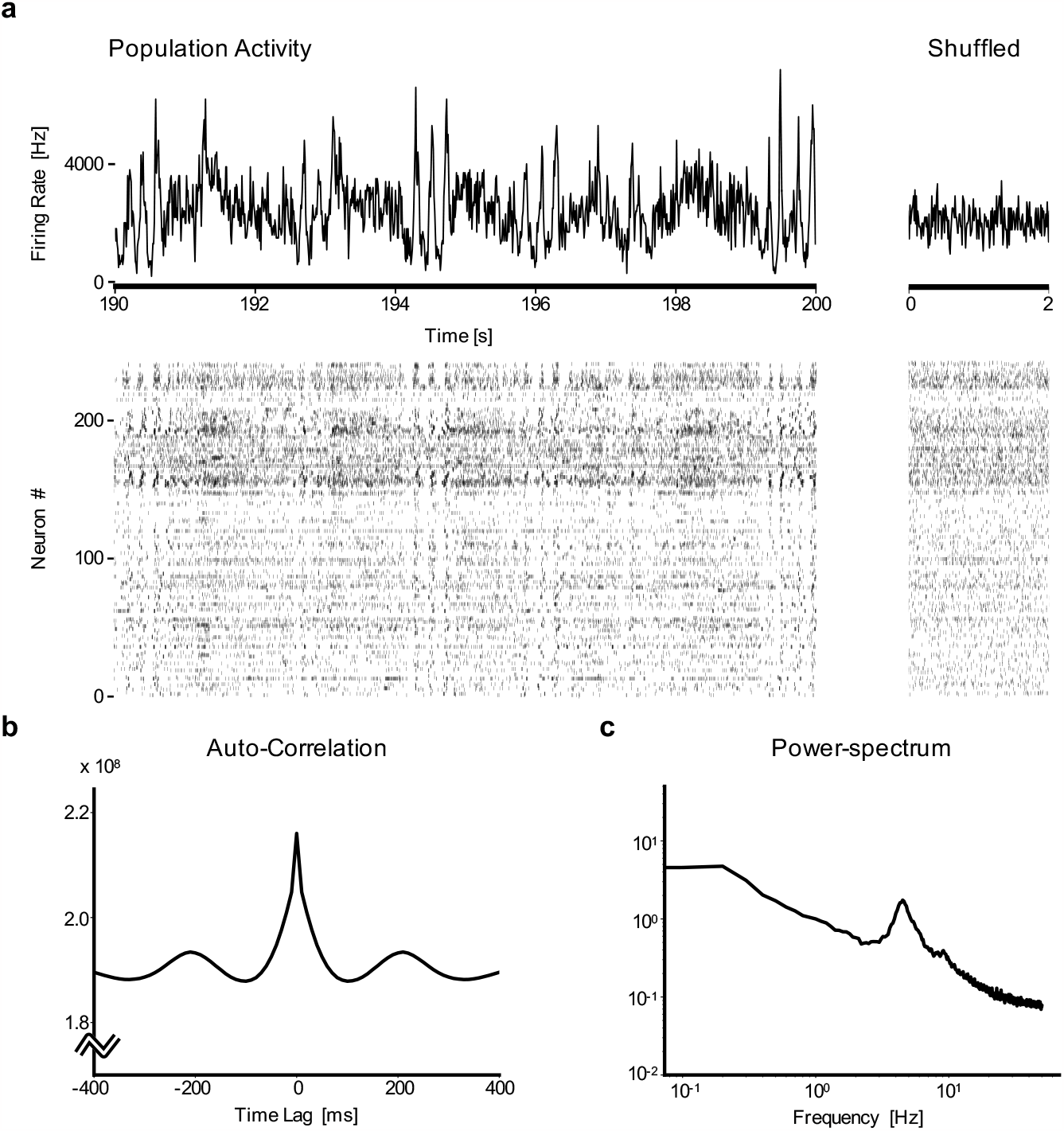
Non-differentiable fluctuations in the population activity of biological neurons. **a** (Left-top) A time histogram of the number of spikes of 242 units for 10 s, with a time bin of 1 ms. (Left-bottom) Spike rasters of 242 units. (Right-top) A time histogram of a shuffled spike train for 2 s. (Right-bottom) Rasters of shuffled spike trains. **b** and **c** An autocorrelation and a power spectrum of the summed spike train.

The autocorrelation of the summed spike train exhibits both slow undulation and a cusp (Fig. 3c). This is reminiscent of our stochastic models (Fig. 2e or 2f), in which a pair of spike trains derived from the rate process in which smooth oscillations and non-differentiable activity are mixed. Because the number of recorded neurons is large and statistical fluctuations are small, we could verify the presence of a sharp cusp at the origin of the autocorrelation.

The power spectrum of an entire spike train exhibits humps with a frequency close to 5 Hz, which may represent the theta rhythm oscillation that is present near the hippocampus (Fig. 3d). The power spectrum exhibits a long tail, which may have contributed to the presence of a cusp in the autocorrelation.

### ShinGLMCC

In the preceding section, we showed that biological data exhibit not only slow undulation but also a cusp at the origin of the CCs. While recent methods for estimating synaptic connections, e.g., GLMCC, have overcome the difficulty of undulation by fitting a smooth function to a given CC (Figs. 1e–g), they do not seem to have successfully captured the cusp in the CC (Figs. 1h–j). This may be because they assumed even smoothness over the time axis of the CC. To further improve the estimate, we need to remove the influence of the cusp.

Here we free the adaptation function from the smoothness at the origin of the time axis. This can be done by removing the penalty at the origin in the prior distribution of the generalized linear model. We call this revised model ShinGLMCC (METHODS).

Although ShinGLMCC has an adaptation function that resembles GLMCC’s, they differ particularly near the origin, whose detail is crucial in determining monosynaptic connections. The adaptation function and synaptic impacts determined by ShinGLMCC are depicted with blue lines in Figs. 1b–j. They seem successful in representing not only undulation but also a cusp of the variation in CCs.

### Comparison of analysis methods on biological data

We applied three analysis methods (Classical CC, GLMCC, and ShinGLMCC) to all pairs of spike trains of 242 neurons recorded from the brain of a rat and obtained connection matrices for all directed links (242 × (242 − 1) = 58, 322).

The Classical CC analysis has suggested 7,103 connections, occupying 12% of all possible links (Fig. 4a). As discussed in the previous section, Classical CC analysis tends to suggest many spurious connections due to large variations in CCs that are found in the biological data.

**FIG. 4.**
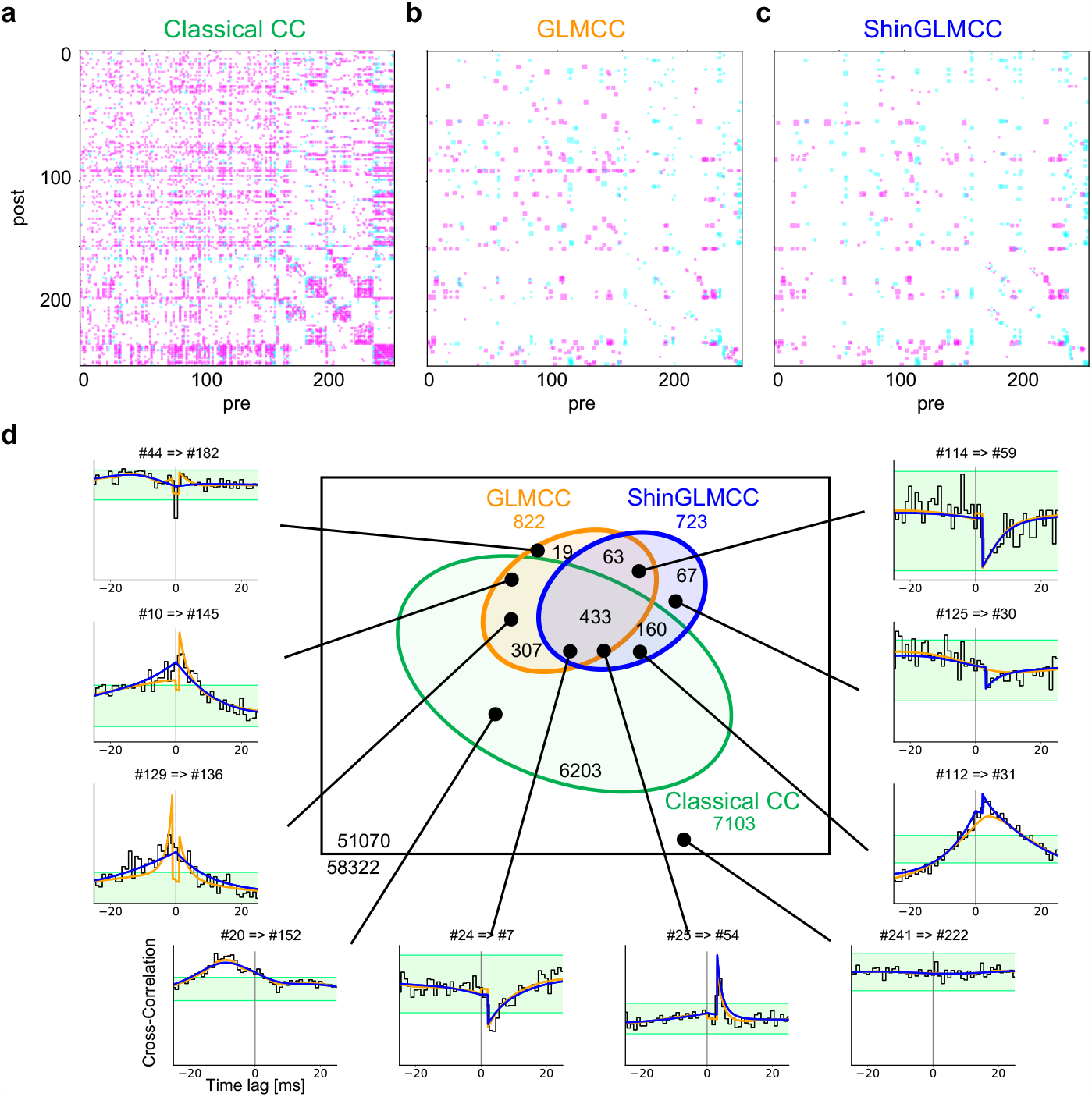
Comparison of analysis methods. **a**–**c** Connection matrices obtained for biological data using the conventional CC method, the GLMCC method, and the newly developed ShinGLMCC method, respectively. Magenta and cyan squares in each matrix represent excitatory and inhibitory connections, respectively estimated by each analysis method. **d** Venn diagram representing the relationships between determined connectivities. Sample CCs are displayed along the periphery.

The GLMCC has suggested 822 connections, occupying 1.4% of possible links (Fig. 4b), obviously improving the inference from Classical CC analysis, by eliminating many seemingly spurious connections.

ShinGLMCC, our new method, suggested 723 connections, occupying 1.2% of possible links (Fig. 4c). A large fraction (69%) of connections suggested by ShinGLMCC are consistent with 60% of connections suggested by GLMCC (Fig. 4c).

The cases for which ShinGLMCC and GLMCC make different suggestions are displayed in Figs. 1h–j and in the periphery of a Venn diagram of Fig. 4d. As has been discussed in the previous sections, ShinGLMCC appears to provide more reliable inferences than GLMCC, particularly for the CCs that exhibit cusps.

### Simulations of a large-scale network of neurons

While inter-neuronal connections determined by the new algorithm ShinGLMCC seem reasonable in the apparent shapes of individual CCs, we cannot be certain of the decision from the available biological data. Here, we compare the performance of Classical CC, GLMCC, and ShinGLMCC in estimating connectivity using synthetic data from a large-scale network of neurons interacting through given synaptic connections.

We performed simulations of networks of 1,000 (800 excitatory and 200 inhibitory) model spiking neurons (METHODS). The mathematical model is similar to what has been used in Ref.[29], but we have explored wider parameter ranges by increasing the strength of interneuronal connectivity such that the system exhibits nonstationary activity.

#### Emergence of non-differentiable activity in large-scale network simulations

With a relatively weak connection strength, the entire system exhibits a stationary time series in which neurons fire asynchronously (Fig. 5a). By increasing the strength of synaptic connections while maintaining a given connectivity matrix, an entire neural network starts to exhibit synchronous burst firings intermittently, displaying irregular stripes in raster diagrams (Fig. 5b). In this strong connectivity regime, a time histogram and raster diagrams resemble those of real biological neurons shown in Figure 3a. By increasing the strength of the connections further, the system starts to exhibit oscillation, in addition to the irregular burst firings (Figs. 5c). The autocorrelation exhibits a cusp and oscillation similar to the biological data as shown in Fig. 3c. This implies that both the smooth oscillation and the non-differentiable fluctuations may emerge even in numerical simulations of a large-scale network of neurons in a regime of strong connectivity.

**FIG. 5.**
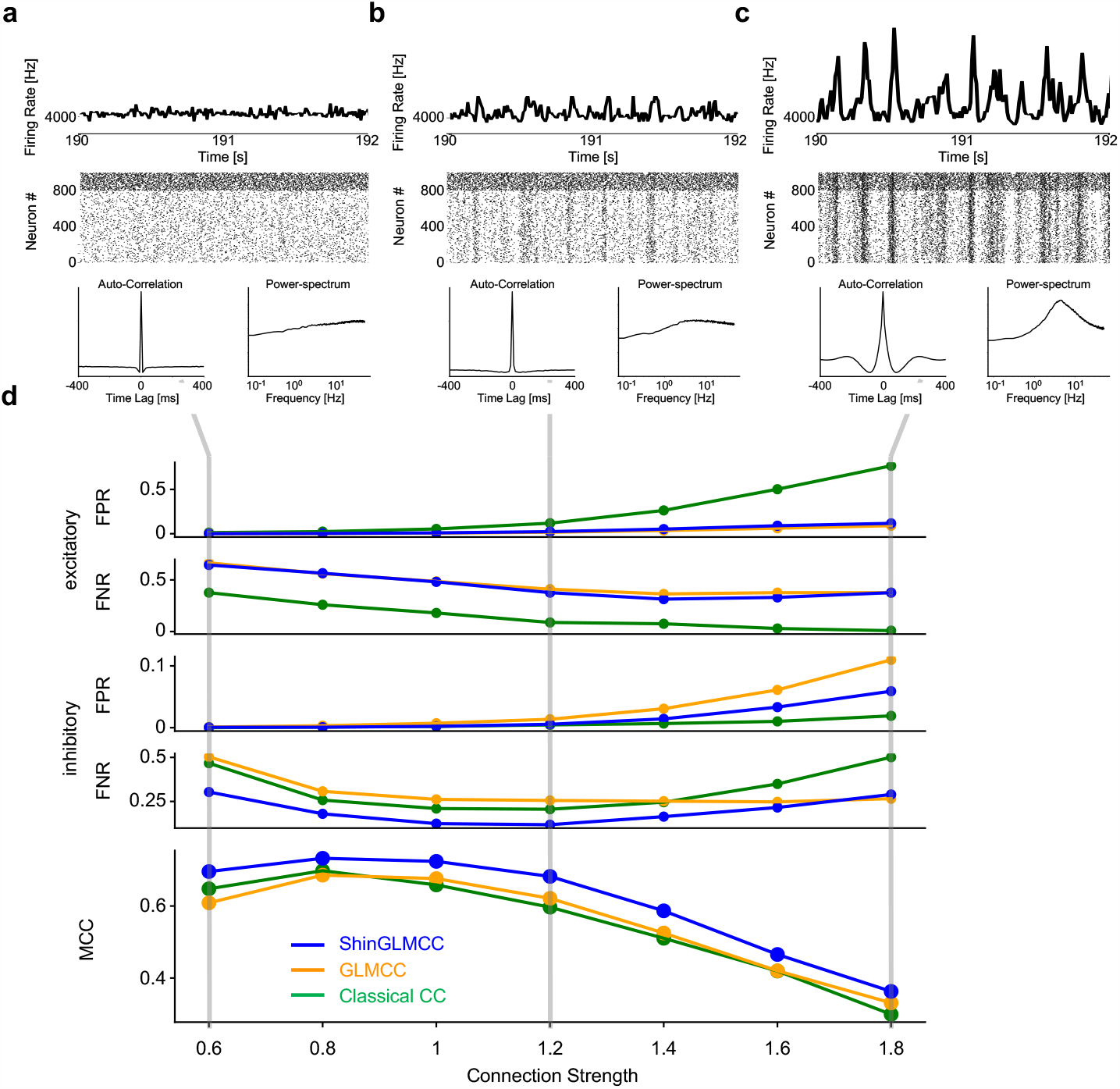
Population activity of a network of 1,000 spiking neuron models. **a**–**c** Time histograms of the number of spikes of a total population of neurons, their spike rasters for a time window of 2 s, autocorrelation functions, and power spectrum of summed spike trains for the simulations of a network of average connection strength *A* = 0.8, 1.2, and 1.6. **d** Estimation performances of Classical CC (green line), GLMCC (orange line), and ShinGLMCC (blue line), as represented by the false positive rate (FPR) and false negative rate (FNR) for excitatory and inhibitory categories computed for 100 neurons (smaller the better), and the Matthews correlation coefficient (MCC) from FPs and FNs (larger the better), plotted against *A*.

#### Evaluation of various analysis methods

We evaluate the ability of Classical CC, GLMCC, and ShinGLMCC to infer connectivity by applying them to 100 spike trains selected from the entire set of 1,000 neurons. We counted the numbers of false positives (FPs) or spurious connections and false negatives (FNs) or missing connections for excitatory and inhibitory categories, and the Matthews correlation coefficient (MCC) evaluated the total performance (METHODS). Fig. 5d shows how the false positive ratio (FPR) and false negative ratio (FNR) for excitatory and inhibitory categories as well as MCC change with the average strength of inter-neuronal connectivity.

The Classical CC analysis registered a large number of FPs for excitatory connections. By contrast, GLMCC and ShinGLMCC performed much better, making fewer FPs or spurious inferences. The performance of ShinGLMCC is better than GLMCC, registering fewer FPs for the inhibitory category than GLMCC, particularly when the network of neurons exhibits synchronous burst firing.

## DISCUSSION

In this study, we have found that CCs of biological spike trains exhibit not only slow undulation but also a cusp at the origin, in addition to possible signs of monosynaptic connectivity. We explored possible mechanisms and found that such undulation and a cusp may have been caused by smooth oscillations and non-differentiable activity in the background, respectively. We developed a new method for estimating connectivity called ShinGLMCC that successfully eliminates the influence of a cusp. We have also performed a simulation of a largescale network of spiking neuron models and found that the model network may also exhibit synchronous burst firings, inducing non-differentiable fluctuations similar to what we have observed in real biological data.

The presence of non-differentiable activity is an inevitable mathematical conclusion derived from the presence of a cusp in a CC. While the concept of non-differentiability cannot be proven rigorously with data of finite recording duration, the biological data analyzed here are consistent with such a mathematical hypothesis. From the cusps observed in CCs of Figs. 1h–j and the cusp in the autocorrelation of a summed spike train in Fig. 3c, we may conclude that the background activity may have been sharp enough to be expressed as “non-differentiable” at least by the timescale of 10 ms or to the frequency of 100 Hz. By drawing our attention to spontaneous activity, we might realize that we have often seen sharp striped patterns in spike rasters of biological data, as shown in Fig. 3b. Accordingly, the concept of non-differentiability may apply to a fairly wide range of biological data.

We also have found that synchronous firings occur in numerical simulations displaying stripes in raster diagrams (Figs. 5a–c) and suggested that these may be related to the non-differentiable activity we found in real brains. Note that a symmetrical peak or a cusp may emerge at the origin of CCs even if there is no nontrivial local connection structure because model neurons are uniformly coupled in a network.

While theoreticians have known that spontaneous coherent dynamics may emerge in large-scale networks and have discussed their possible functions in information processing [30–38], they have not paid attention to the sharp time profile of coherent synchronous firings in a way we described as non-differentiability in the present study. Sharp fluctuations in the entire network may imply that the system may respond quickly to a given stimulus, possessing a high computational capacity. It is interesting to consider the conditions for networks of excitable elements exhibiting non-differentiable activity and their possible functions for information processing.

We also have devised fast algorithms for the estimation methods GLMCC and ShinGLMCC and provided user-friendly software packages for researchers to obtain connection diagrams and individual analysis details for all neuron pairs from a given set of recorded spike trains. The algorithm of ShinGLMCC finishes computation for 58,322 pairs in about 30 min on a Mac Pro (2019, 28cores).

While the recorded neurons are still a mere small portion of the entire neuronal network, the disclosure ratio is rapidly increasing. The number of neuronal pairs that should be examined for potential connectivity increases much more rapidly with the order of the square of the number of neurons. With a suitable and fast analysis tool of this kind, we will be able to explore the difference in neuronal circuitry across different functional regions in the brain.

## METHODS

### Cross-correlogram (CC)

The CC is a means to display the potential interdependence of two units with a time delay *t*, defined as

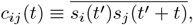

where 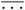 represents an average over *t*^*′*^, *s*_*i*_(*t*) and *s*_*j*_(*t*) are spike trains of *i*th and *j*th neurons, respectively (Fig. 1a). For a pair of independent spike trains, *c*_*ij*_(*t*) is generally flat, being accompanied by sample fluctuations (Fig. 1b). The classical theory instructed that *c*_*ij*_(*t*) may exhibit humps or dips at several milliseconds from the origin if two neurons are interacting via excitatory or inhibitory synapses, respectively (Figs. 1c and 1d), and devised a method of detecting them as an evidence of data outlying a given statistical significance level [17, 40].

However, CC may exhibit large fluctuations even if two neurons are not directly connected, because they may be receiving common background activity in the brain. We have seen that many CCs obtained from biological spike trains exhibit large undulation and a cusp (Figs. 1e– j). Hence, the Classical CC analysis often fails in its estimation.

### Model consideration of various variations in CCs

To determine possible mechanisms by which CCs exhibit undulation and a cusp, we ran inhomogeneous Poisson processes, such that spikes are derived from given instantaneous rates. Their rates should contain coherent parts so that the CC exhibits large deviations that exceed normal statistical fluctuations.

#### Models producing smooth undulation in a CC

The smooth undulation observed in a CC may be produced by simulating two neurons whose firing rates are modulated with smooth oscillations, as represented by

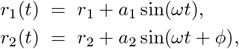

where *ω* is the frequency of the underlying oscillation and *ϕ* is the phase lag (Figs. 2a and 2b). The CC for the spike trains derived from the above-mentioned two processes can be obtained analytically as

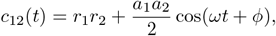

thus reproducing the slow undulation. Note that this is a theoretical result obtained for spike trains of infinite length. CCs constructed from a finite number of spikes exhibit noisy fluctuations. Nevertheless, such fluctuations can be minimized if we can obtain spike trains of sufficiently long duration.

The slow undulation appearing in biological CCs generally exhibits damping because the underlying oscillation is not purely harmonic. For the cases in which frequencies are distributed normally around *ω* with variance 1*/δ*^2^, the CC exhibits damping, as represented by the Gabor function,

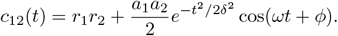

While many biological CCs show a peak at the origin, there are cases in which the peak is shifted from the origin. Thus two neurons exhibiting a peak shifted from the origin of the CC may have been receiving brainwave oscillations with a fixed latency.

#### Models that produce a cusp in a CC

Even if the undulation of a biological CC does not seem simply harmonic, any smooth shape can be represented as a sum of many oscillatory components. Accordingly, undulations of nontrivial shape may also be explained such that the neurons may have been influenced by many brainwave components. However, the cusp we found in many biological CCs cannot be explained as a simple combination of a finite number of smooth brainwave components. Instead, the original processes should be “mean square differentiable” [41]. We have explored mathematical models that can produce a cusp at the origin of the CC. For simplicity’s sake, we assume that a pair of neurons generate spikes according to identical time-dependent rates. While their rate processes may contain independent parts in practice, they should have common inputs of this kind.

In the first model, the firing rates of two neurons follow the Ornstein-Uhlenbeck process [42], as given by the stochastic differential equation,

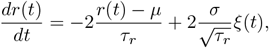

where *μ, σ*, and *τ*_*r*_ are the mean, standard deviation, and the time scale of the fluctuation. *ξ*(*t*) is Gaussian white noise characterized by ensemble averages ⟨*ξ*(*t*) ⟩ = 0 and ⟨*ξ*(*t*)*ξ*(*t*^*′*^)⟩ = *δ*(*t* − *t*^*′*^). The rate is continuous but is non-differentiable everywhere (Fig. 2c). In this case, the CC of two neurons generating spikes according to the identical rate processes *r*(*t*) can be obtained analytically as (Fig. 2c)

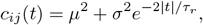

exhibiting a cusp at the origin [43–45]. These two neurons are assumed to have no synaptic connections because they are generating spikes independently according to instantaneous rates. The cusp appeared with zero time lag because the rate processes are coherently driven.

In the second model, the firing rates of two neurons follow a switching rate process in which the rate changes discontinuously between high and low values

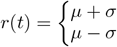

according to the “Markov switching process” or “random telegraph process” with a mean time interval *τ*_*r*_ [45–49] (Fig. 2d). In this case, the inter-transition interval *t* is distributed exponentially as 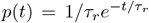. In this case, the CC of two neurons generating spikes becomes 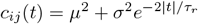, which is identical to the CC of the Ornstein-Uhlenbeck process.

In the Ornstein-Uhlenbeck process, the time-dependent rate is non-differentiable everywhere. In the Markov transition process, the rate changes discontinuously at a finite rate. It is noteworthy that such different processes may give rise to an identical cusp in CCs. A common characteristic of the underlying rate processes for producing a cusp in the CC is that the firing rates of two neurons undergo coherent non-differentiable rates.

The original rate process cannot be identified uniquely from a CC, because a CC is obtained by averaging over time series. Even though we cannot visualize the original rate processes from recorded spike trains, we can obtain a sharp profile of the CC from sufficiently long spike sequences because the number of spike counts in each bin of the CC increases with the length of the spike trains.

### Methods of estimating monosynaptic connectivity

#### GLMCC

Because Classical CC analysis has made many false inferences due to large undulations in CCs, modern analysis methods attempted to eliminate the influence of the undulations. For instance, the GLMCC method [28] introduced a function *a*(*t*) to absorb the large undulation in an original CC and tried to fit the function

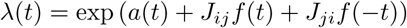

to *c*_*ij*_(*t*) obtained for a given pair of neurons. Here *J*_*ij*_ represents a discrete impact due to possible monosynaptic connectivity from the *j*th neuron to the *i*th neuron, and *f* (*t*) is the time profile of the synaptic interaction, modeled as 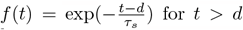 and *f* (*t*) = 0 otherwise, where *τ*_*s*_ is a typical timescale of synaptic impact and *d* is the transmission delay. For making a fitting function *a*(*t*) represent a smooth undulation in the CC, a large gradient of *a*(*t*) is penalized with the prior distribution

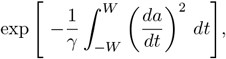

where *W* is the time window of a CC and *γ* is a hyperparameter representing the flatness of *a*(*t*); we selected here as *W* = 50 ms and *γ* = 5 × 10^*−*4^ [ms^*−*1^]. The set of parameters {*J*_12_, *J*_21_, *a*(*t*) } was determined by maximizing the posterior distribution given the spike data {*t*_*k*_} in the CCs [28] or its logarithm:

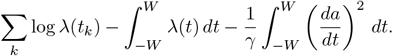

#### ShinGLMCC

While the original GLMCC algorithm solved the problem of undulation by fitting a smooth function evenly to the original CC, it cannot reproduce a cusp at the origin. Therefore, we have revised it into ShinGLMCC by allowing the function *a*(*t*) to bend at the origin. Specifically, we separate a single integral in the prior distribution into two pieces of half-lines and changed the first-order derivative to the second-order derivative,

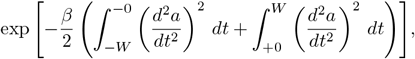

with the constraint that the value *a*(*t*) is continuous at the origin.

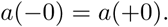

This means that *a*(*t*) is requested to be smooth in each half region, except at the origin. Because the order of derivative changed, we have selected a new penalizing parameter of different dimensionality, *β* = 10^6^ [ms^*−*3^]. We have devised a numerical code enabling fast estimation of ShinGLMCC.

While the new fitting function *a*(*t*) optimized by the new ShinGLMCC may look similar to the one optimized by the original GLMCC, they can be different particularly near the origin, deriving different decisions for detecting connections in some cases as demonstrated in Figs. 1h–j.

#### Likelihood ratio test

In the original GLMCC, the presence of connectivity was determined by thresholding the estimated connectivity parameter |*J*_*ij*_| depending on the recording time, and firing rates of pre- and post-synaptic neurons. However, this thresholding causes an asymmetry in detectability between excitatory and inhibitory connections. Accordingly, GLMCC was revised by introducing the likelihood ratio test such that the connection is determined based on the likelihood ratio between the presence and absence of the connectivity,

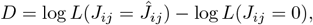

where 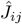 is the estimated connection parameter and *L* is the model likelihood [29]. Based on Wilks’ theorem [50], we may reject a null hypothesis that a connection is absent (or determine that the connection is likely to present) if 2*D > z*_*α*_, where *z*_*α*_ is the threshold of *χ*^2^ distribution of a significance level *α*.

The ShinGLMCC algorithm also adopts the likelihood ratio test with *α* = 10^*−*4^. We also searched for the timescale of synaptic impact *τ*_*s*_ out of 1, 2, 3, and 4 ms, and the transmission delay *d* out of 1, 2, and 3 ms, based on the likelihood ratio.

### Evaluating analysis methods

The performance of estimation methods was evaluated using the data obtained from simulations of a large-scale network of neurons. For this purpose, we have counted the number of estimation failures such as FPs and FNs for excitatory and inhibitory categories (for the excitatory category, we considered only EPSP *>* 1 mV). These failures are quantified by false positive rate (FPR) and false negative rate (FNR) defined as,

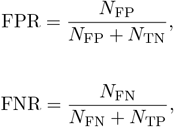

where *N*_FP_, *N*_TN_, *N*_FN_, and *N*_TP_ represent the numbers of false positive, true negative, false negative, and true positive connections, respectively. The smaller, the better.

As a total score of the goodness of estimation, we computed the Matthews correlation coefficient (MCC) [51] defined by

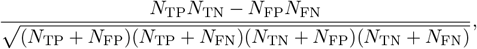

the larger, the better. We have taken the macro-average MCC that gives equal importance to excitatory and inhibitory categories,

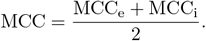

### Simulating a large scale network of neurons

To evaluate estimation methods such as Classical CC, GLMCC, and ShinGLMCC in their accuracy in detecting connectivity, we have generated spike trains from 1,000 model neurons interacting with fixed synaptic connectivity. While the model simulation is similar to that of Ref.[29], we carried out an independent simulation with the following conditions.

As a spiking neuron model, we adopted the Multitimescale Adaptive Threshold (MAT) model [52, 53]. In this model, “membrane potential” *v*_*m*_ obeys a simple leaky integration of input signal without resetting:

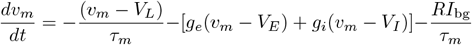

where *g*_*e*_ and *g*_*i*_ represent the excitatory and inhibitory conductances, respectively, whereas *RI*_bg_ represents the background inputs coming from outside the population. The conductance evolves with

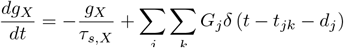

where *X* stands for excitatory *e* or inhibitory *i τ*_*s,X*_ is the decay constant, *t*_*jk*_ is the *k*th spike time of *j*th neuron, *d*_*j*_ is a synaptic delay and *G*_*j*_ is the synaptic weight from *j*th neuron. *δ*(*t*) is the Dirac delta function.

In this model, the threshold for generating spikes *θ* (*t*) is lifted when generating a spike and subsequently decays with two time-scales as

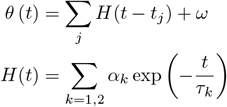

where *t*_*j*_ is the *j*th spike time of a neuron, *ω* is the resting value of the threshold, *τ*_*k*_ is the *k*th time constant, and *α*_*k*_ is the weight of the *k*th component (*k* = 1, 2). The shorter timescale *τ*_1_ = 10 ms represents the membrane time constant, and the longer timescale *τ*_2_ = 200 ms represents the adaptation of neuronal spiking. The parameter values chosen for the present simulation are summarized in Table I.

**TABLE 1.**
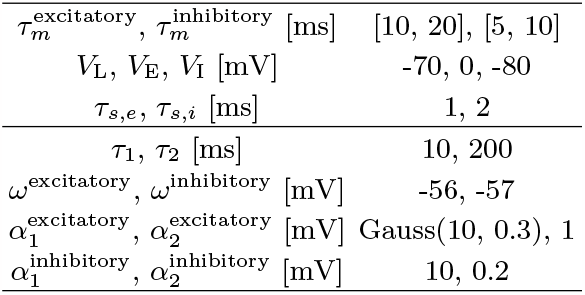
Parameters for neuron models

Among 1,000 neurons, 800 excitatory neurons innervate to 12.5 % of other neurons with EPSPs that are log-normally distributed [54–56], whereas 200 inhibitory neurons innervate randomly to 25 % of other neurons with IPSPs that are normally distributed.

The excitatory and inhibitory synaptic connections were sampled from the respective distributions. The excitatory conductances 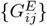 were sampled from a lognormal distribution with the mean −5.543 and SD 1.30 of the natural logarithm of the conductances [54, 55]. The inhibitory conductances 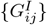 were sampled from the normal distribution with the mean 0.0217 and SD of 0.00171. If the value sampled from the normal distribution is negative, it is resampled from the same distribution. The entire connection matrix {*G*_*ij*_ } is multiplied by the strength constant *A*, which we changed from 0.6 to 1.8 to see if the network exhibits nonstationary burst firing.

The synaptic delays from excitatory neurons are distributed uniformly between 3 and 5 ms, and those from inhibitory neurons are distributed uniformly between 2 and 4 ms.

We added a background current to represent random bombardments from excitatory and inhibitory neurons of the outside population [57].

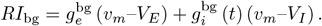

The dynamics of the conductances can be approximated as a stationary fluctuating process represented by the Ornstein–Uhlenbeck process,

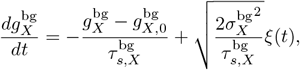

where *g*_*X*_ stands for *g*_*e*_ or *g*_*i*_, and *ξ*(*t*) is the white Gaussian noise satisfying ⟨*ξ*(*t*) ⟩ = 0 and ⟨*ξ*(*t*)*ξ*(*s*) ⟩ = *δ*(*t* − *s*). The parameter values for the background inputs are summarized in Table II.

**TABLE II.** Parameters for background inputs.

|  |  |
| --- | --- |
| $\tau_{s,e}^{\text{bg}}, \tau_{s,i}^{\text{bg}}$ [ms] | 2.7, 10.5 |
| $g_{e,0}^{\text{bg}}, g_{i,0}^{\text{bg}}$ | 1.85, 4.83 |
| $\sigma_e^{\text{bg}}, \sigma_i^{\text{bg}}$ | 0.245, 0.400 |

The numerical simulation of a given network was carried out for a time interval of 3,600 s with a time step of 0.0001s.

### Code and data availability

We have developed the ShinGLMCC algorithm using Python. We also revised the GLMCC algorithm so that the computation can be carried out much faster than the original one developed in Refs. [28, 29]. Additionally, we developed the network simulation code in Python. All the code is available at the repository, https://github.com/yasuhirotsubo/neuroscience such that connection matrices estimated by ShinGLMCC and GLMCC can be computed for a given set of spike trains.

### Biological data

We have analyzed publicly sourced data of spike trains recorded in parallel from the brain of a freely moving rat using a silicon probe called Neuropixels [39]. They recorded multiple neurons from the visual cortex, hippocampus, and thalamus in an awake mouse. We analyzed spike trains of 242 units according to their evaluation as being well-isolated.

## Acknowledgements

We would like to thank the research groups of Ref. [39] who have provided their experimental data open to the public. We also thank Hiroya Nakao for providing the mathematical knowledge regarding the mean square differentiability, and Peter Kirkwood for constructive comments on the manuscript. Y.T. was supported by JSPS KAKENHI 22K12186 and JSPS KAKENHI 22H05511. S.S. was supported by JSPS KAKENHI 22H05163 and by New Energy and Industrial Technology Development Organization (NEDO).

## Author contributions

Y.T. and S.S. designed the study and developed the analysis method. S.S. drafted the manuscript and performed theoretical analysis. Y.T. developed codes, analyzed experimental data, and performed numerical simulations.

